# Evolutionary analysis of gene ages across TADs associates chromatin topology with whole genome duplications

**DOI:** 10.1101/2021.06.11.448047

**Authors:** Caelinn James, Marco Trevisan-Herraz, David Juan, Daniel Rico

**Author notes:** Joint first authors (Caelinn James, Marco Trevisan-Herraz). Lead contact and corresponding author: Daniel Rico (, @danielrico_bio).

## Abstract

Topologically associated domains (TADs) are interaction sub-networks of chromosomal regions in 3D genomes. TAD boundaries frequently coincide with genome breaks while boundary deletion is under negative selection, suggesting that TADs may facilitate genome rearrangements and evolution. We show that genes co-localise by evolutionary age in humans and mice, resulting in TADs having different proportions of younger and older genes. We observe a major transition in the age co-localisation patterns between the genes born during vertebrate whole genome duplications (WGDs) or before, and those born afterwards. We also find that genes recently duplicated in primates and rodents are more frequently essential when they are located in old-enriched TADs and interact with genes that last duplicated during the WGD. Therefore, the evolutionary relevance of recent genes may increase when located in TADs with established regulatory networks. Our data suggests that TADs could play a role in organising ancestral functions and evolutionary novelty.

**Highlights:**

- TADs coincide with clusters of genes that are close in their evolutionary age.
- Whole genome duplications mark a transition in gene age co-localisation clusters.
- Gene age co-localisation patterns are associated with TAD insulation.
- Young essential genes share TADs and interact with old genes.

## Introduction

Recent advances in understanding the modular organisation of chromatin are beginning to shed light on the local context within which genes are situated, thanks to the development of chromosome conformation capture (3C) techniques.^1,2^ In 3C data, we can see that chromosomes are composed of sets of neighbouring genes and regulatory regions that tend to preferentially interact with each other in three-dimensional (3D) space, called topologically associated domains (TADs).

Accumulating data suggests that TAD boundaries could amount to important functional elements, providing a modular system that might enhance the evolvability of genomes. Recent work has shown that synteny breakpoints between different mammals^3,4^ and flies^5^ are enriched at TAD boundaries. Certain classes of paralog protein-coding genes have been shown to co-localise within TADs and partially share regulatory elements in mammals^6,7^ and clusters of conserved noncoding elements of key developmental genes coincide with TADs in humans and flies^8^. TAD boundaries are evolutionarily constrained^9^ and their deletion is under negative selection.^10,11^ Interestingly, a comparison of linkage disequilibrium (LD) blocks and TADs revealed that TAD boundaries do not coincide with recombination hotspots, suggesting that recombination might be deleterious at these regions.^12^ Taken together, these results indicate that TAD boundaries are conserved but breakable,^13^ facilitating genome rearrangements during evolution while preserving intra-TAD functional and regulatory interactions. They also support the Integrative Breakage Model of genome evolution,^14^ where chromatin conformation facilitates genome reorganisation during evolution.

Evolution is not just driven by rearrangements of the 3D structure of the genome; gene duplication events also play a role in evolution. Most gene duplication events affect genomic regions containing one or a handful of genes, but whole chromosome and whole genome duplications (WGD) can also occur and be important drivers of evolution.^22^ There has been at least one WGD event (but most likely two) in the ancestor of vertebrates, and the majority of genes duplicated during this event present in extant vertebrates have not duplicated since ^23–25^. More recently, it has been shown that the expansion of many gene families after the WGDs was frequently associated with gene specialisation evidenced by restricted expression and recruitment of novel, tissue-specific regulatory elements contributing to the increase in the regulatory complexity in vertebrates.^27^

Gene duplication events (whether individual genes or WGD) can allow us to determine the evolutionary “age” of a gene based on its most recent duplication event.^20^. The function, expression patterns, and protein-protein interactions of genes are associated with their age, where older genes more frequently perform essential functions, are expressed in more cell types, and show more interactions.^15–17^ Evolutionarily old genes are more likely to be essential genes,^17–19^ but we still do not understand the role genome architecture may play in the acquisition of essential functions.

Here, we investigated whether protein-coding genes that share a TAD had also originated around the same evolutionary time (and thus share the same evolutionary age). We show how gene age is non-randomly distributed in the human and mouse genomes, where genes within the same TAD tend to have similar gene ages. This gene age co-localisation is lost in those TADs containing more inter-TAD interactions in the 3D genome; these TADs also tend to be enriched in young genes. TADs enriched in genes born before or during the WGDs tend to co-localise in TADs that rarely contain younger genes, but when they do, the young genes are more likely to be essential than expected by chance and show more 3D chromatin interactions with older essential genes. Overall, we uncover an intimate relationship between genome architecture and gene evolution, with implications for the emergence of essential functions and the evolution of regulatory circuits in evolutionarily young genes.

## Results

### Gene ages are not randomly distributed in human and mouse TADs

There are two main ways to define the age of a gene. First, we can define how old a gene is by looking for homologues in other species (with sequence similarity searches such as BLAST) and defining the gene age by the last common ancestor of the species where homologues are found **(Figure 1A)**. In reality, this approach is helpful in estimating the age of the gene family: the age of the root of the phylogenetic gene tree.^20^ To estimate the age of each individual gene within a gene family, we can consider when was the last time that the gene was involved in a gene birth event **(Figure 1B)**,^21^ where the most common mechanism is gene duplication. In this definition, both parental and daughter genes share the age of the corresponding duplication event.

**Figure 1.**
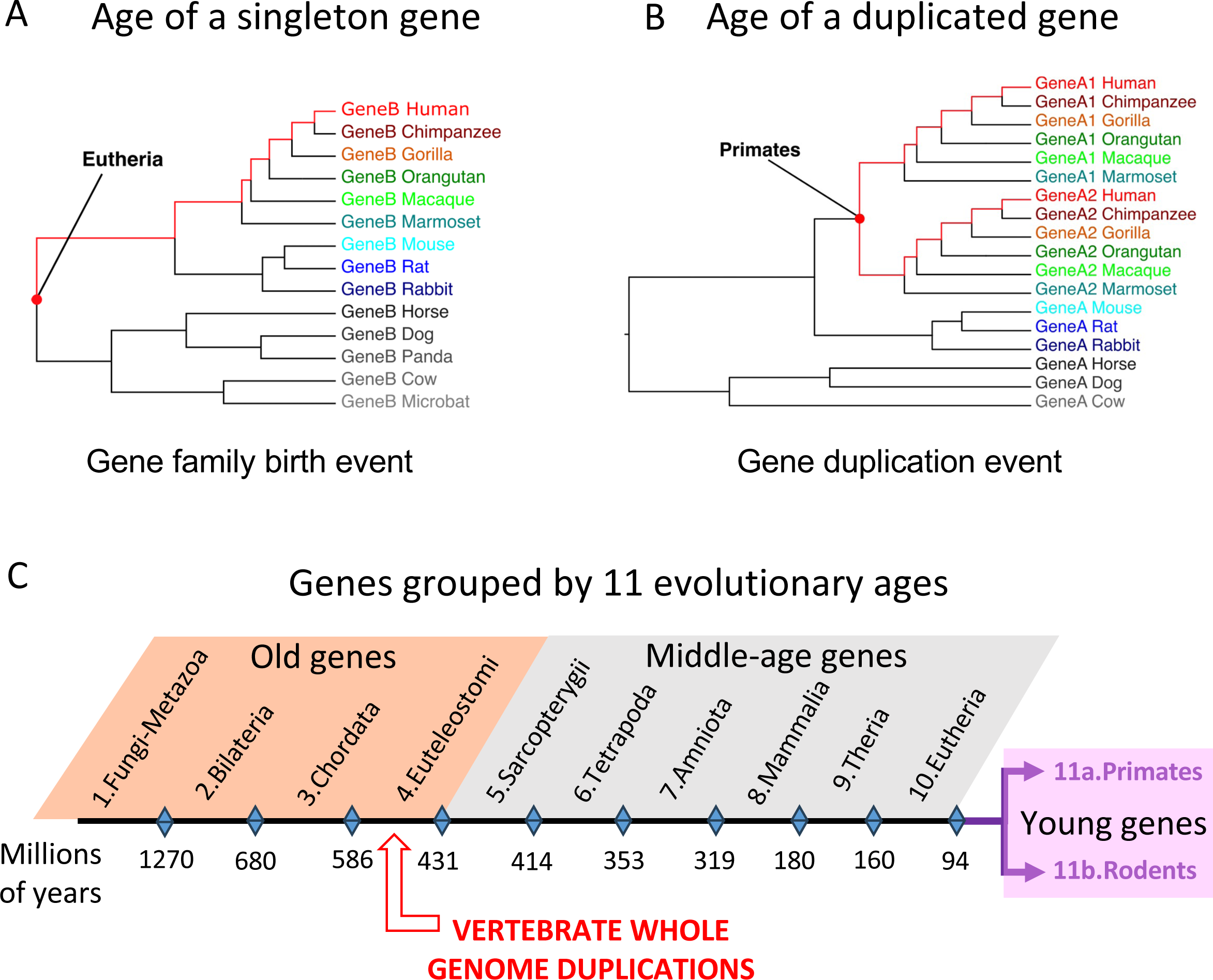
Diagrams showing how gene ages are dated. **A.** Toy gene tree illustrating how the gene age corresponds to the last common ancestor for singleton genes. **B.** Toy gene tree illustrating how the gene age corresponds to the most recent duplication node for duplicated genes. As for singleton genes, the gene age indicates the last time a gene was involved in a gene birth event. **C.** Human and mouse genes were grouped in 11 gene age groups (see **Table 1** for more details).

Human and mouse protein-coding genes were classified into 11 age groups based on each gene’s most recent gene birth event^29,30^, using gene trees retrieved from Ensembl^28^ (**Figure 1C** and **Table 1**). Ten age groups correspond to shared ancestors between humans and mice (from FungiMetazoa to Eutheria) while the youngest age group in each species corresponds to Primate (in humans) or Rodent (in mice) gene births; see **Table 1** and **STAR Methods** for more details.

**Table 1.**
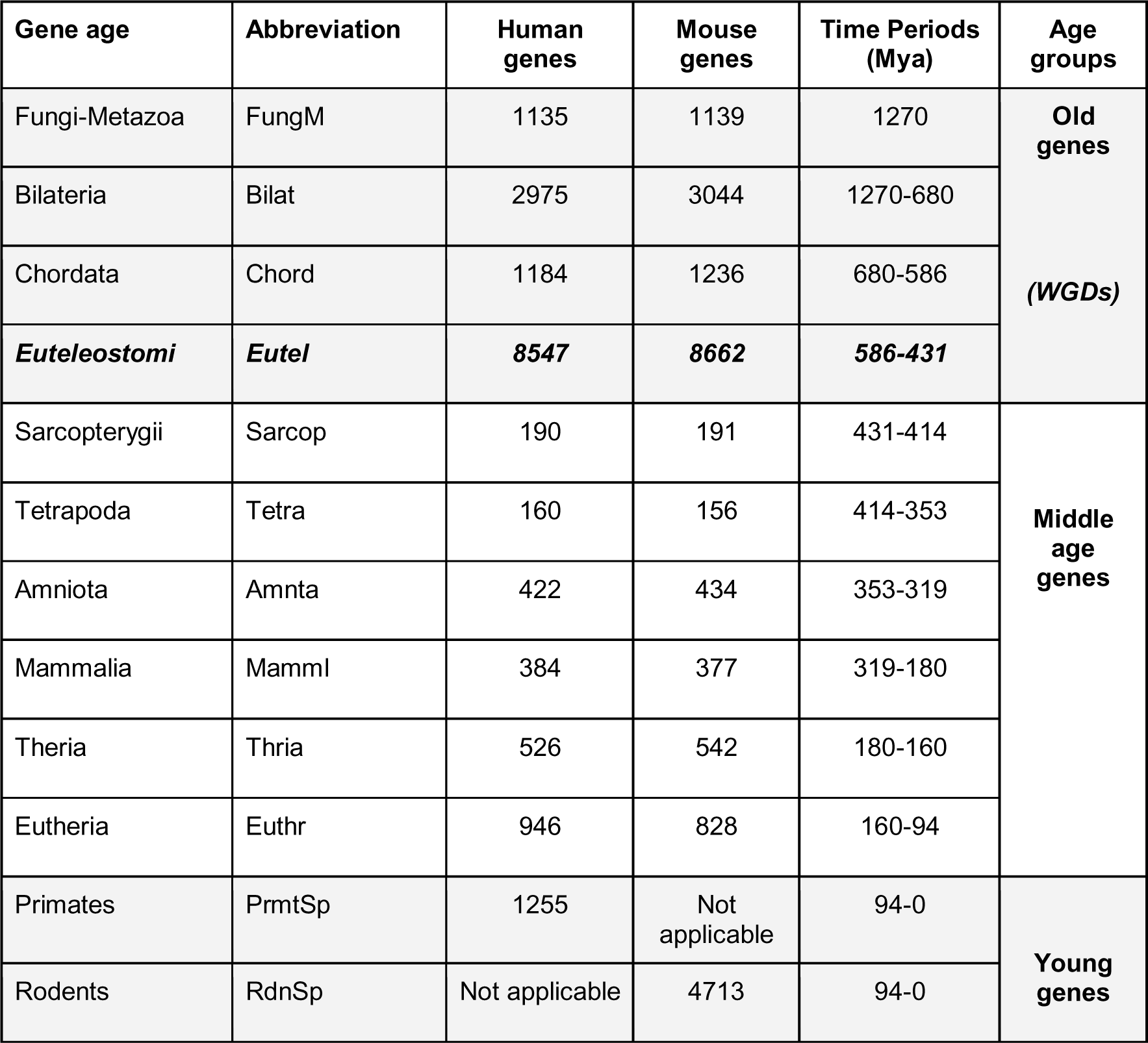
Gene ages used in this study, including the abbreviations used in some figures, number of human and mouse genes in each age group, and the evolutionary time periods covered (in millions of years). Euteleostomi genes correspond to those born during the WGDs. All ages are shared between human and mouse, except Primates (Primates specific, or PrmtSp, in human) and Rodents (Rodent specific, or RdnSp, in mouse), see **STAR Methods** and **Key resources table** for details.

For each age group in humans, we computed the frequencies of finding genes of every other age in the same TAD, using TAD maps available from human embryonic stem cells (hESCs, **Key resources table**) .^31^ We used data from ESCs because they perform some of the oldest functional processes in eukaryotic evolution, such as symmetric and asymmetric cell division, and they constitute themselves one of the evolutionarily older cell types in multicellular organisms.^32–34^ To test the significance of this age co-localisation, we calculated the p-values for the probability of finding gene pairs having each combination of ages (i.e. the probability of obtaining a gene of age *i* in the same TAD by chance when selecting a gene of age *j*). To calculate these p-values, we considered the null hypothesis to be the distribution of these probabilities when randomising the ages of the genes (and hence across TADs, see **Figure 2A**, **Figure S1** and **STAR Methods**). To visualise the p-values, we generated a matrix showing the probability of finding each combination of gene ages (**Figure 2B**).

**Figure 2.**
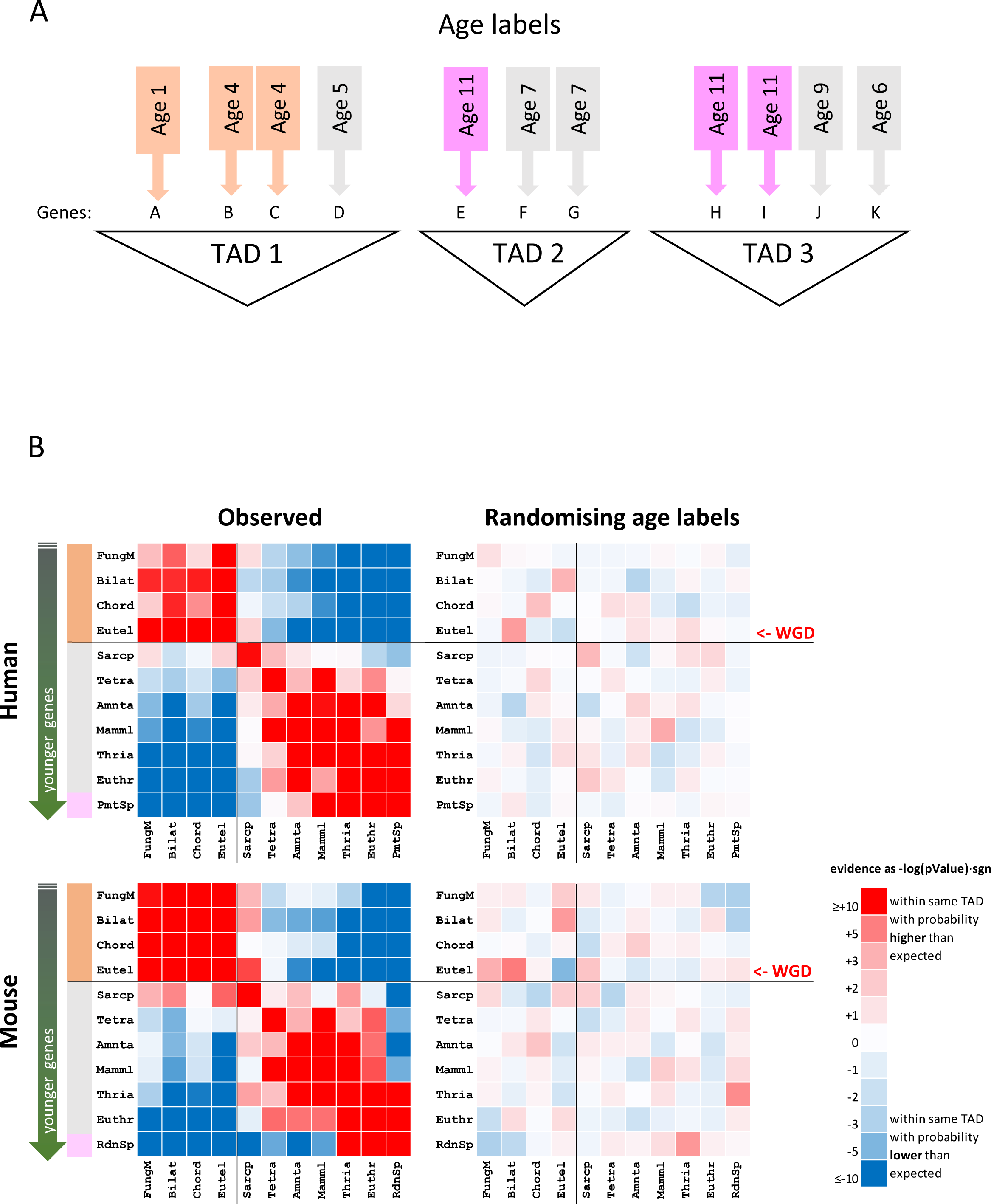
Genes co-localise by age in TADs. **A.** Cartoon representing three TADs and their genes. In the co-localisation analysis, genes are kept in their original locations and gene age labels are randomly relocated. **B.** Representation of the matrices displaying the evidence that genes with any combination of two ages are found together in the same TAD by using the natural cologarithm of the p-value, using a positive (negative) sign when this is higher (lower) than expected. Increasing intensities of red (blue) show increasing evidence that the combination is found together more often (less often) than expected by chance, white representing that the match found is exactly as expected by chance. See **Table 1** for the correspondence between abbreviated and full gene ages. The colour legend next to the abbreviated age names indicate the three gene age groups depicted in Figure 1C; the gene age corresponding to the whole genome duplication (WGD) is indicated with black horizontal and vertical lines. **Left:** observed p-values for human and mouse. **Right:** representative results of the same operation after randomising 100 times the age labels of each gene (so each age group keeps the same number of genes, but just assigned randomly).

The age co-localisation results show that gene ages are not distributed randomly in the TADs of the human genome (**Figure 2B**, **left**, and **Figure S2**). For the sake of simplicity, we merged gene ages into three major groups: old (genes not duplicated since the WGDs with Euteleostomi or older ages), middle-age (from Sarcopterygii to Eutheria) and young genes (Primates, or Rodents in the case of mouse; see **Figure 1C** for details). Surprisingly, our analysis shows a clear co-localisation pattern according to two gene age groups. The old genes are significantly co-localised within the same TADs. However, they share TADs with middle-aged and young genes less often than expected by chance. In contrast, we see significant TAD co-localisation of young and middle-aged groups (**Figure 2B**).

To further test the observed pattern of gene age co-localisation, we randomised the age labels of each gene and repeated the analysis. We observed no age co-localisation when age labels were randomly assigned **(Figure 2B**, **right**, and **Figure S3)**. This suggests that the co-localisation effect we saw between old gene ages (FungiMetazoa to Euteleostomi) on the one hand, and middle-aged to young genes on the other hand, was not random, but instead an effect of genome structure.

To assess whether this result is ESC-specific or depends on the TAD definitions used (which can vary depending on the TAD-calling method),^1,35^ we performed the same analysis using independently defined TAD maps from different cell types: human B-cell derived lymphoblastoid GM12878 cells (derived from the highest resolution HiC data in humans to date),^36^ monocytes, neutrophils and naive CD4 T-cells^37^ (**Key resources table**). In all these cases, very similar two age co-localisation groups were also observed (**Figures S4** and **S5**).

Within the group of old genes (see **Figure 1C**), Euteleostomi genes (or ohnologs) are the most recent, corresponding to gene duplications that resulted from WGDs that occurred at the origin of the ancestral vertebrate genome.^23–25^ If the marked gene age co-localisation we observed in humans is associated with the gene birth events before and after the WGDs, we would expect to see a similar pattern in the mouse genome (as they share all the gene ages until the Primates-Rodents split).

Therefore, we performed the same analysis using mouse genes and TADs from mouse embryonic stem cells (mESCs, derived from the highest resolution HiC data in mouse to date^38^ , **Key resources table**). Interestingly, a very similar significant pattern was observed in the mouse (**Figure 2B**, **bottom-left**) that disappeared when the age labels were randomised (**Figure 2B**, **bottom-right**). It is particularly remarkable that Primate and Rodent genes show the same pattern, although they represent independent gene birth events.

In summary, the results in both species show that there is a major transition in the TAD co-localisation patterns prior to Sarcopterygii, coinciding with the time when the two rounds of WGD are thought to have occurred.^25^

### Gene age co-localisation is stronger in TADs than in fixed-size partitions of the human genome

As genes within the same TAD are close to each other, we need to consider that the observed co-localisation of gene ages could be TAD-independent. To test this alternative hypothesis, we divided the human genome into windows of fixed size, using different sizes (ranging from 20 Kb to 50 Mb), and repeated the analysis for each window size (**Figure 3**). For each window size, we generated a corresponding matrix of p-values and generated a heatmap. A visual comparison of the results obtained suggests that fixed windows in the size range of TADs (300 Kb to 1Mb) show similar patterns of co-localisation, but are less evident than real TADs (**Figure 3**).

**Figure 3.**
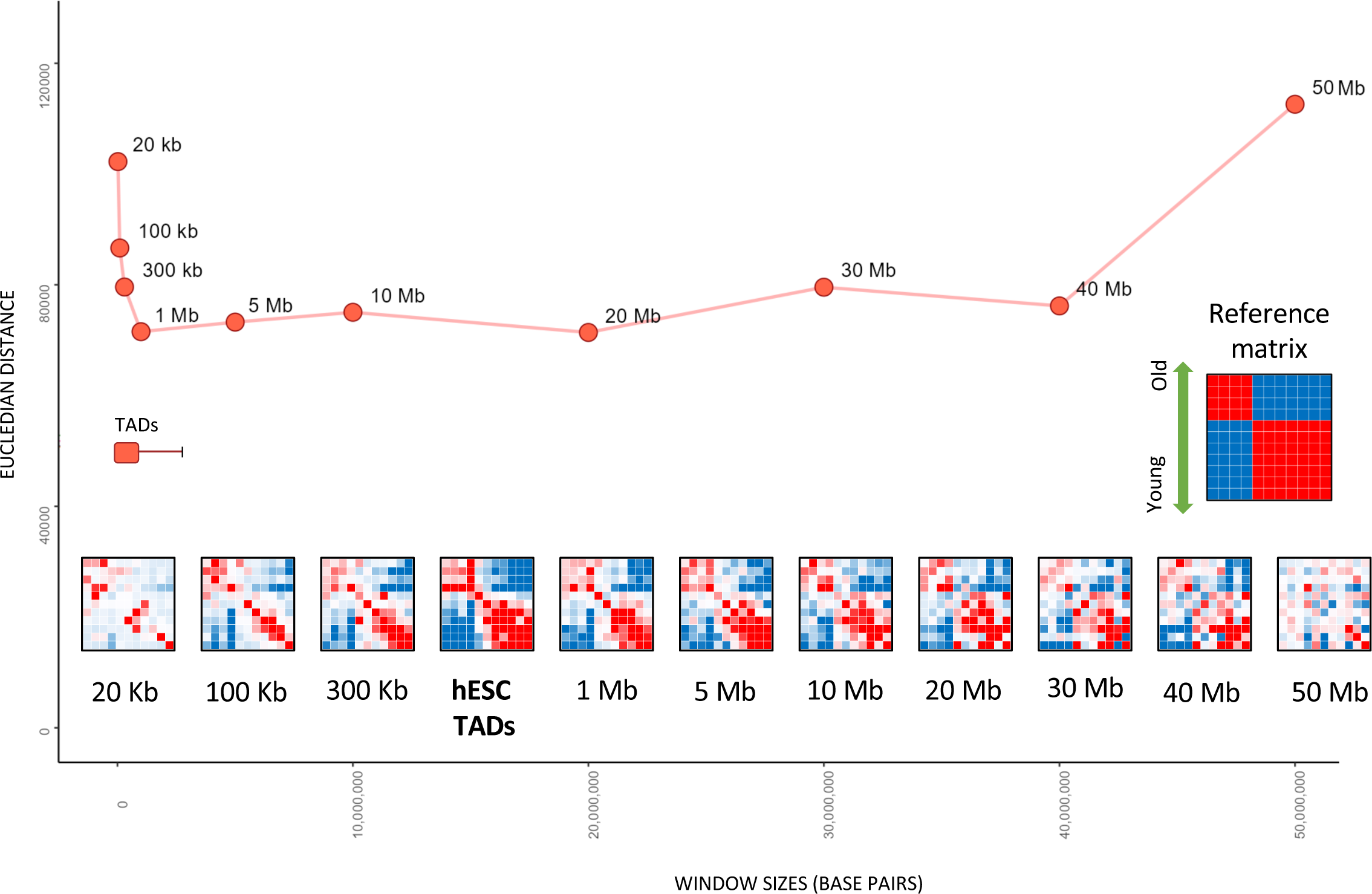
Comparison of co-localisation of genes of different ages in both TADs and fixed-size windows. Each heatmap at the bottom represents the gene age co-localisation p-values (see colour legend and more details in Figure 2) obtained using the fixed sized windows of the indicated sizes - from 20 Kb to 50 Mb. The heatmap with p-values obtained with the hESC TADs (same as in Figure 2) is included for comparison. The points in the graph show the quantitative comparison of the p-value matrices with a reference matrix (top right) that represents an extreme two-group co-localisation pattern. The y-axis indicates the Euclidean distance of each real p-value matrix to this reference. The x-axis represents the size of the fixed-size window used (and the ranges of sizes for TADs).

To quantitatively compare the results of different windows and real TADs, we calculated the similarity (Euclidean distance) of each observed matrix of p-values to an extreme pattern of co-localisation where genes perfectly cluster in either the four older (FungiMetazoa-Euteleostomi) or seven younger (Sarcopterigii-Primates) age groups. The Euclidean distance between two numeric matrices is a measure of the straight-line distance between their respective points in the multi-dimensional space defined by the matrices’ rows or columns (**STAR Methods**). This approach showed that the matrix with co-localisation p-values obtained with the TADs has a shorter Euclidean distance (meaning a higher similarity) than any fixed-size windows, indicating that the co-localisation pattern is stronger in TADs. Therefore, the observed results cannot be simply explained by the linear proximity of genes.

We found that the 20Kb and 50Mb windows had the highest Eucledian distances. In the case of the 50Mb window, the window covers the whole chromosome for some of the shorter chromosomes and so the co-localisation pattern that emerges is more likely representing the co-localisation of gene ages by chromosome. In the case of the 20Kb window, the window is so small that it does not capture many genes at once, thus does not generate many meaningful comparisons.

### Gene age co-localisation patterns are related to TAD insulation

TAD boundaries can show different degrees of insulation^39^ : the higher the insulation, the fewer inter-TAD interactions are observed (**Figure 4A**). Boundaries with higher insulation are generally associated with more CTCF binding^40,41^ and are thought to prevent undesired enhancer-promoter interactions between different neighbouring regulatory programmes.^42^ To further investigate the connection of gene age co-localisation with TAD structure, we studied its relation to the insulation of TADs. For this, we used the TAD boundary insulation scores calculated by Gong and co-workers^43^ using the high-resolution HiC data^36^ of the human lymphoblastoid cell line GM12878 (**Key resources table**). We calculated a TAD insulation score by averaging the insulation scores of their boundaries and then divided the TADs into five quintiles based on their insulation (from very low to very high, **Figure 4B**). We found that the gene age co-localisation pattern was different depending on the TAD insulation strength, which was almost lost in the two groups with the lowest insulation strength (**Figure 4B**).

**Figure 4.**
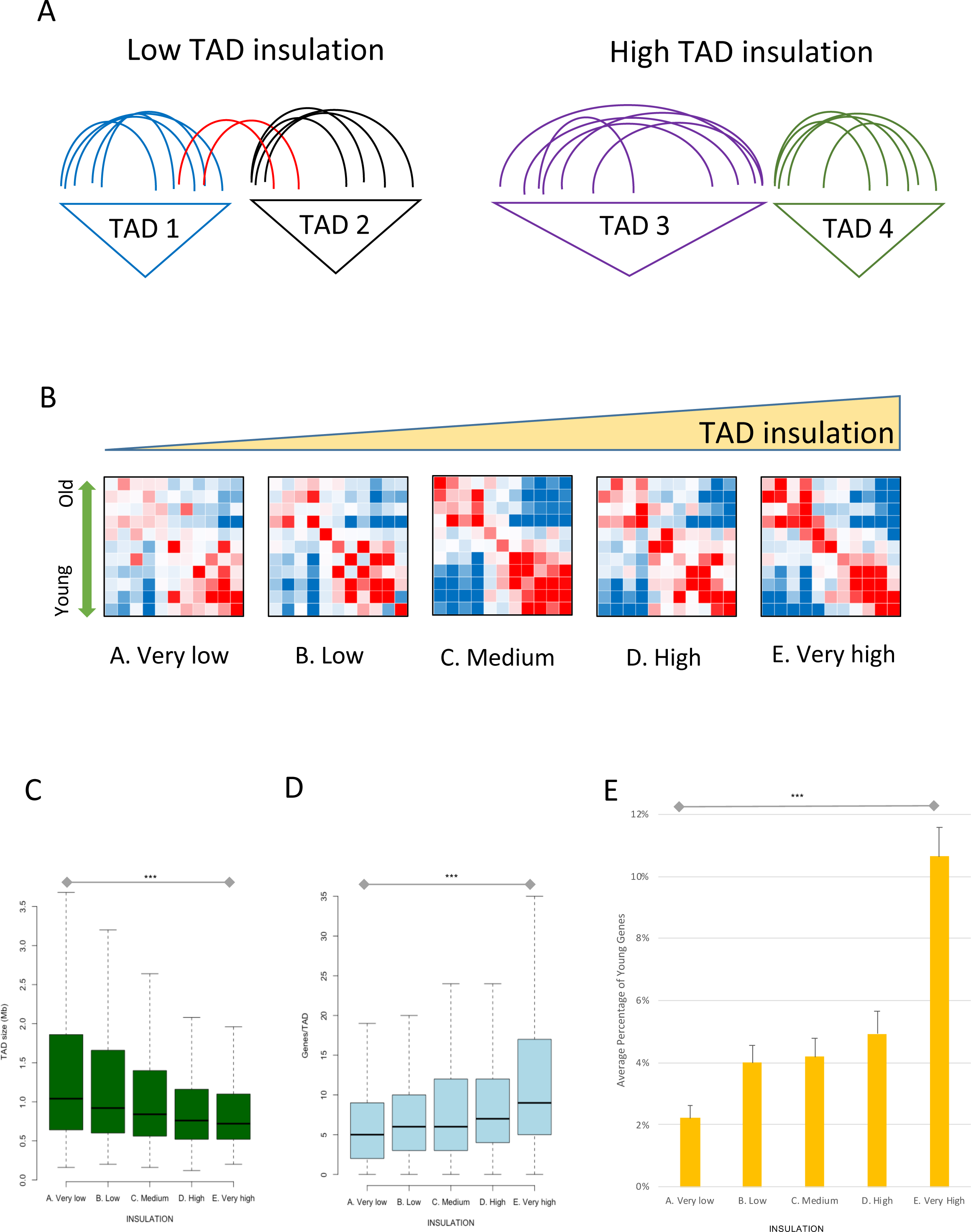
TAD boundary insulation is associated with gene age co-localisation patterns. **A.** Cartoon illustrating TAD boundary insulation; the more inter-TAD 3D interactions that occur, the lower the insulation strength between two neighbouring TADs. **B.** Co-localisation p-values obtained for the five groups of TADs of GM12878 lymphoblastoid cells, based on their boundary insulation strength. For details see Figure 2 and legend. **C.** TAD size distributions for the five insulation groups of TADs in panel B. **D.** Distributions of the number of genes per TAD insulation group. Error bars indicate the standard errors. **E.** Percentage of Young (Primates) Genes for the five TAD insulation groups. Significance of Wilcoxon’s (**C, D**) and chi-squared (**E**) tests: *** indicates p < 0.001.

Interestingly, the strongest two-group pattern was observed in those TADs with intermediate insulation. High and (especially) very high show a stronger co-localisation pattern near the diagonal showing a higher resolution of the gene age co-localization and suggesting different gene birth dynamics.

The gene age co-localisation patterns can also be affected by other factors that might vary among these five groups of TADs, such as the size of the TADs or their number of genes. Therefore, we decided to investigate the characteristics of TADs with differing insulation strengths. We observed that more insulated TADs tend to be significantly smaller (p = 4.33 x 10^-12^, comparison between the very low and very high insulated groups, **Figure 4C**) but have a significantly higher gene density (p < 2.2 x 10^-16^, **Figure 4D**). Next, we analysed how the youngest genes born during the Primates lineage varied among the TAD insulation groups: Young genes are relatively more enriched in more insulated TADs (p < 2.2 x 10^-16^, **Figure 4D**). In summary, these results show that the more insulated TADs are, the more gene density they have in general, but they also have a relatively higher proportion of young genes. The combination of these properties probably contributes to the differences in gene age co-localisation patterns associated with the TAD insulation strength.

TAD boundaries can be stable or conserved across many cell types, or restricted to one or a few cell types. By analysing the TADs in 37 human cell types, it has been shown that the TAD boundaries that are more stable across cell types are associated with higher evolutionary constraints, CTCF binding, and housekeeping genes.^9^ Therefore, to further study the observed conservation of the co-localization pattern in different cell types, we investigated if human TAD stability^9^ (**Key resources table**) was associated to different proportions of gene ages. We consider TADs to be stable if their boundaries are shared among at least 5 cell types, while the boundaries of unstable TADs are shared by 4 cell types or less, as this threshold splits the distribution into two groups of similar size (see **STAR Methods**). We compared the ratio of Old-depleted TADs (those that contained less than 10% of old genes) between stable and unstable TADs and found that unstable TADs have a significantly higher ratio of Old-depleted TADs (**Figure S6**). In other words, TADs unusually rich in young (Primate) and middle-age genes tend to be less stable across cell types.

### Essential young genes are associated with TADs enriched in old genes

Evolutionarily old genes are more frequently expressed across cell types and are more often classified as “essential” than younger genes ^18,19^. Essential genes are required for growth, development and reproduction, either at the cellular level or at the whole organism level ^18,44^, and loss of function of these genes can compromise the viability or fitness of the organism ^18,19^. It is assumed that the reason that evolutionarily older genes are more likely to be essential is due to older non-essential genes gradually being removed by evolution.

However, it is still unclear how some young genes become essential. We therefore chose to investigate the differences in TAD environments between essential young genes and non-essential young genes, to understand whether TAD location may influence the probability of young genes becoming essential.

We examined how the proportion of essential genes in a TAD varied depending on the ages of the genes within the TAD. We used the database of Online GEne Essentiality^44^ (OGEE, **Key resources table**) to determine whether each gene was essential or not. As expected, as a TAD increases in the proportion of young genes, it decreases in the proportion of essential genes, whilst the opposite was true for the proportion of old genes **(Figure 5A)**.

**Figure 5.**
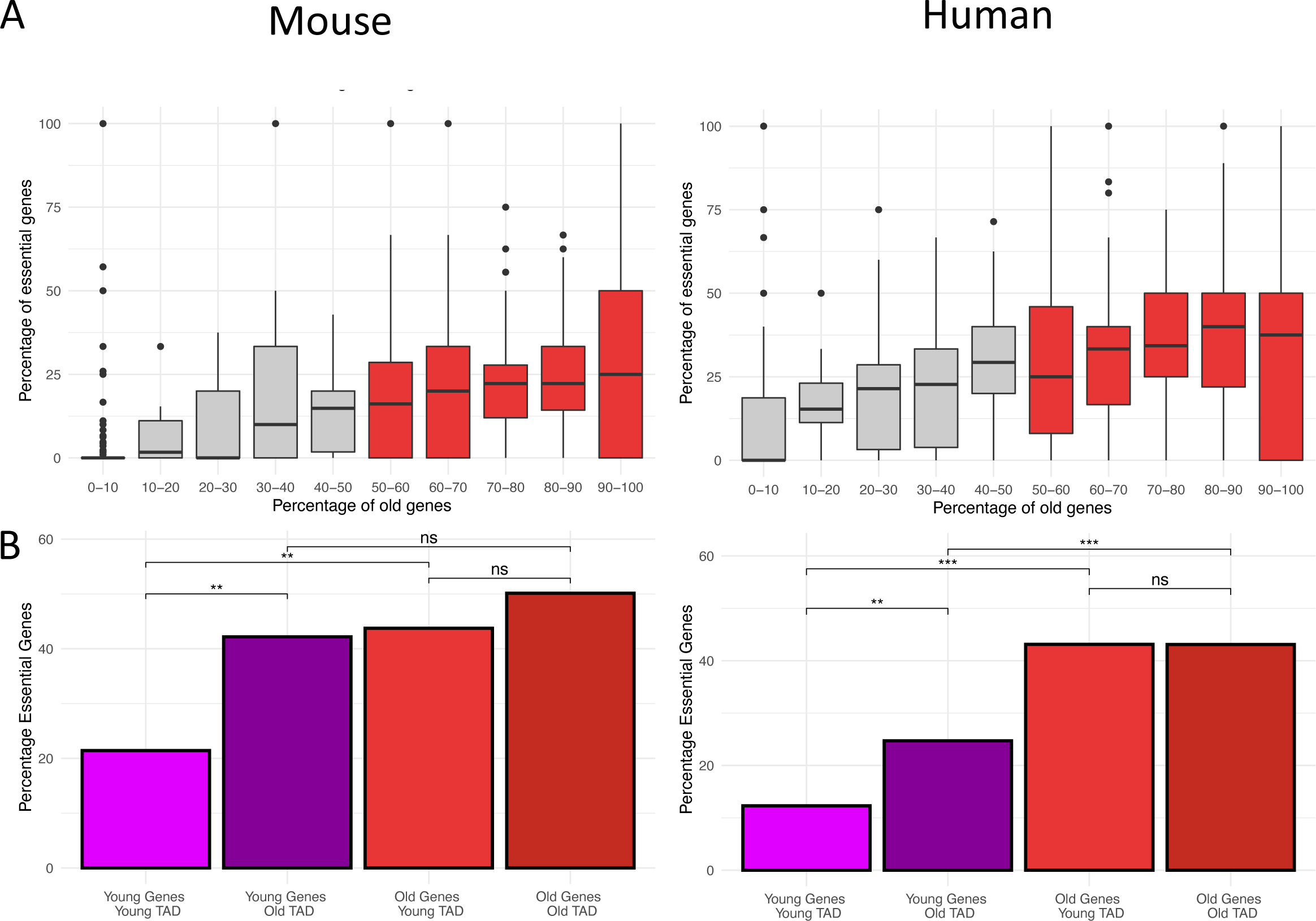
Different proportions of young genes, old genes and essential genes across TADs in mice and humans. **A.** Variation of the percentage of essential genes in comparison to the percentage of old genes. Left: mouse; right: human. **B.** Percentage of essential genes across four gene ‘groups’ - young genes in Young-enriched TADs, young genes in Old-enriched TADs, old genes in Young-enriched TADs and old genes in Old-enriched TADs. Left: mouse; right: human.The asterisks indicate the p-values when carrying out chi-squared tests between the different groups. Significance of chi-squared tests: * indicates p < 0.05, ** indicates p < 0.01, *** indicates p < 0.001, (ns: non significant).

Next, we investigated if young essential genes were enriched in TADs with other young genes or with older ones. For this, we focused our analyses on two types of TADs: ‘Old-enriched TADs’, in which at least 50% of the genes were old (Euteleostomi or older, **Figure 1C**); and ‘Young-enriched TADs’, in which at least 50% of the genes were Primate (in human) or Rodent (in mouse). We grouped the genes in these TADs depending on whether they were young genes in Young-enriched TADs, young genes in Old-enriched TADs, old genes in Young-enriched TADs, and old genes in Old-enriched TADs, and calculated the proportion of genes in each group that were essential (**Figure 5B)**. We then carried out chi-squared tests between each group to determine if there were any significant differences.

We found in both the mouse and the human dataset that young genes in Young-enriched TADs were less likely to be essential than young genes in Old-enriched TADs (humans: p = 5.54 x 10^-3^, mice: p = 1.29 x 10^-3^), whilst the frequency of essential old genes was similar in Young-enriched and Old-enriched TADs (humans: p = 1, mice: p = 4.49 x 10^-2^). These data suggest that the age of neighbouring genes can help determine whether or not a young gene is essential or functionally important, while for old genes, the age of the neighbouring genes does not seem to be informative (**Figure 5B**).

### Essential young genes show more 3D interactions with old genes and other essential genes

The enrichment of essential young genes in Old-enriched TADs could be associated with its incorporation with evolutionary older regulatory networks, which would result in an increased number of chromatin 3D interactions with older genes. To explore this hypothesis, we analysed publicly available high-resolution promoter-capture HiC (PCHi-C) networks from mouse^45^ and human cells^37^ (**Key resources table**). Each network consists of nodes that represent the PCHi-C fragments and edges that represent their interactions.^46^ To compare the interactions of young essential genes with those of young non-essential genes, we constructed a sub-network from the PCHi-C data in each species, keeping only interactions that involve nodes with lineage-specific genes (**STAR Methods**). From there, the subnetwork was divided into two further subnetworks: one subnetwork where every pair of interacting nodes contained a node with an essential young gene, and a second subnetwork where every pair of interacting nodes contained a node with a non-essential young gene. For both subnetworks, we looked at the age and essentiality of the interactors (**Figure 6A**).

**Figure 6.**
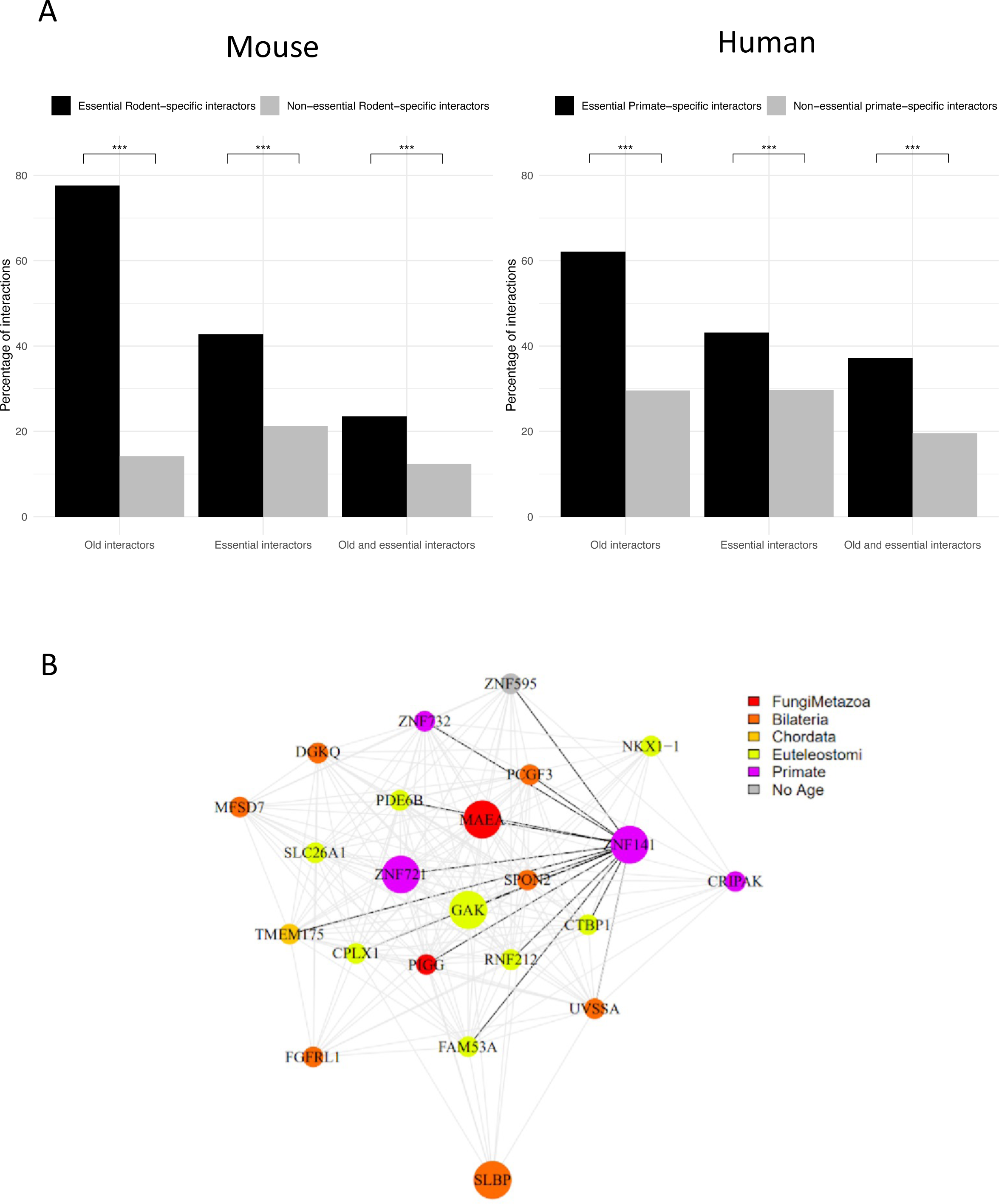
Young essential genes interact more frequently with old and essential genes. **A.** Comparison of PCHiC interactions with old, essential, and old and essential interactors for essential Rodent and non-essential Rodent genes in mESCs (left) and essential Primate and non-essential Primate genes in human CD8 T-cells (right). Asterisks indicate that p-values for the chi squared tests were all below 0.001. **B.** HiC interaction network for the TAD containing ZNF141. Nodes represent genes within the TAD, whilst the edges represent the interactions between the genes. Essential genes are represented with a large node, whilst non-essential genes are represented with a small node. Black edges represent interactions involving ZNF141. The age of a gene is represented by the colour of the node.

We found that essential young genes have more chromatin interactions with old genes than non-essential young genes in both mice (p = 7.7 x 10^-90^) and humans (p = 9.2 x 10^-21^). Essential Rodent and Primate genes also showed significantly higher interactions with other essential genes, and a similar trend was observed when we looked at interactors that were both old and essential (**Figure 6A**). These results suggest that, independently of any definition of TADs, 3D interactions with old and/or essential genes are associated with a higher probability of a young gene being essential. Although we do not know how many of these physical interactions are actually regulatory, these enrichments suggest that young essential genes have a higher probability of being co-regulated with old and/or essential genes.

Zinc Finger Protein 141 (ZNF141) is a representative example of a young human essential gene (based on CRISPR-Cas9 screening data)^44^ located in a TAD rich in old genes; out of the 27 genes in the TAD, 22 (81.5%) are aged Eutelostomi or older (**Figure 6B**). This TAD is in the telomeric region on chromosome 4p16.3 and contains another 4 essential genes, three of which are directly interacting with ZNF141 in hESC DNAse HiC data^31^: MAEA (Bilateria), GAK (Euteleostomi) and ZNF721 (Primates). Interestingly, mutations in ZNF141 have been shown to be associated with postaxial polydactyly type A malformation characterised by well-formed functionally developed 5th digit duplication in hands and/or feet.^47^

## Discussion

### Genes co-localise in the genome by age

Previous works have shown that synteny breakpoints in mammals^3,4^ and flies^5^ tend to be located near TAD boundaries. Although we still do not know what are the underlying causes that explain these observed correlations, it is tempting to speculate that regulatory or functional interactions within some TADs might be important to maintain. Indeed, other independent works have reported that TAD boundaries show evolutionary constraints.^9,10,11^ Here, we have found that the TADs in human and mouse genomes coincide with clusters of genes that are close in their evolutionary age. The local expansion of duplicating gene families is probably a key factor contributing to the co-localisation of younger genes. It is possible that the structure of TADs that are rich in old genes is under negative selection, and the birth of new genes there is detrimental. This might have been important at least right after the WGDs and then promoted by the local expansions of genes afterwards, which tend to accumulate in error-prone late replicating regions of the genome.^29^ Of course, this observation does not imply any causative relationship between the WGDs and TAD structure, as many other factors could have contributed to these gene age co-localisation patterns. However, our results suggest this is an intriguing hypothesis that deserves to be investigated in the future.

Ohnolog pairs have been shown to exhibit higher spatial proximity in 3D nuclear organisation than other paralog pairs.^48^ Taken together with our own data, these observations suggest that the constraints of the WGD could have influenced the genome architecture in mammalian genomes. Our results comparing the co-localisation of gene ages in TADs and windows of fixed sizes indicate that the TAD structure may potentially be playing a role in the observed gene age distribution. These results show the gene age co-localisation is stronger when using TADs than when we used fixed sized windows of different sizes, suggesting that there is a significant association between TAD locations and gene age distribution in the genome. As the main difference we observe is between genes born during the WGD (Euteleostomi) or earlier, we can speculate that TAD structure could have helped to maintain regulatory interactions among genes within old-gene rich TADs and buffer, to some extent, the effect of regulatory interactions with younger genes. However, it is important to note that we only explored chromatin physical interactions, and future perturbation experiments will be needed to determine which ones are actually regulatory.

Another possibility is that the lower rates of mutations and copy number variants in early replicating TADs, rich in old genes, reduce the probability of genes being born there, independently of negative selection biases. Early replicating TADs are expected to have more efficient DNA repair^49^ to reduce the probability of having damaging mutations; the low frequency of young genes in these TADs there might be a secondary effect of a lower mutation rate in these regions. Future availability of TADs in a wider diversity of species will allow to investigate the conservation of these TADs and their chromatin interactions.^50–52^

Interestingly, we noticed that gene age co-localisation patterns vary depending on TAD insulation and the co-localisation is actually not observed in lowly insulated TADs. In contrast, very high insulated TADs are more enriched in recently born genes and genes there tend to co-localise more with genes with genes born immediately before or after them. This could reflect a more dynamic evolutionary history in these TADs, an interpretation that is compatible with the higher gene density that we have observed on them. We also found that more insulated TADs tend to be smaller despite showing higher gene density, suggesting that it might be more complicated to insulate larger regions. A recent unpublished comparative analysis of TADs in different 4 primates and 4 rodents identified ultraconserved TADs shared by humans and the other 7 species.^52^ Interestingly, these ultraconserved TADs show higher insulation strength and contain genes with cell type specific patterns.

### The essentiality of genes varies by gene age

The enrichment of young essential genes in old-enriched TADs could be associated to different factors: the higher probability of the chromatin in these TADs to be open, functional relationships with the same essential processes where their neighbours are involved or new regulatory and/or structural roles keeping the integrity of the 3D structure. Indeed, the potential ability of TADs to encapsulate regulatory environments might allow them to preserve and protect them from the rise of other younger environments.

Some of these young essential genes could indeed have originated from parental genes that were already essential in the ancestral species; in this case, the essentiality itself would be older than the gene age. Moreover, enhancers have been shown to be a possible substrate for the origin of new genes^53–55^ and it is also possible that the essentiality retrieved from CRISPR screens is actually capturing the essentiality of the regulatory role of the genomic regions. Indeed, promoters with enhancer function (ePromoters) of interferon signalling response have been shown to be more conserved than other non-enhancer genes belonging to the same pathway.^56^

## Conclusion

Ultimately, our findings have a simple but very important implication: TADs can be classified by the age of their genes. This can be useful to study the evolution of different biological processes, such as regulatory modules of development, cell type ontogenies or cancer.

Here, we explored the relationship between TADs, gene age and gene essentiality, showing that young essential genes tend to be located in TADs rich in genes that are older than them. This could help in prioritising the study of genes of unknown function by taking into account the age of their neighbours.

### Limitations of the Study

The estimation of gene ages of human and mouse genes depends on the quantity and variable quality of species genome assemblies and gene annotations available, as well as the quality of automatic gene tree reconstructions. Initiatives such as the Earth Biogenome Project^57^ will help to have better species coverage and finer resolution of gene ages. However, this will come with additional computational challenges associated with gene tree reconstructions.

This study is also limited to the HiC-derived TAD datasets from humans and mice that were generated by different laboratories and processed in different ways. Analysing more TAD data from HiC uniformly processed and harmonised for different cell types across more mammalian species, together with the availability of age data for the different species in the future, will enable us to explore whether the results are consistent across more cell types and species.^50–52^

## Author contributions

Conceptualization: CJ, MT, DJ and DR; Analysis: CJ, MT and DR; Writing: CJ, MT, DJ and DR; Supervision, DJ and DR; Funding Acquisition, DR.

## Acknowledgements

We are grateful to Maninder Heer and Maria Rigau for their critical comments. We thank Prof Aristotelis Tsirigos for sharing their TAD insulation data. This work was supported by a Wellcome Trust (https://wellcome.ac.uk) Seed Award in Science (206103/Z/17/Z) awarded to DR.

## STAR Methods

### RESOURCE AVAILABILITY

#### Lead contact

Further information and requests for resources should be directed to and will be fulfilled by the lead contact, Daniel Rico.

#### Materials availability

There are no newly generated materials associated with the paper.

#### Data and code availability

- This paper analyses existing, publicly available data. These accession numbers for the datasets are listed in the **Key resources table**.
- All original code has been deposited at https://github.com/ricolab/TAD_Evolution and https://doi.org/10.5281/zenodo.10067329 and is publicly available as of the date of publication. DOIs are listed in the **Key resources table**.
- Any additional information required to reanalyse the data reported in this paper is available from the lead contact upon request.

### METHOD DETAILS

#### Dating ages in genes

An age was assigned to mouse and human genes using Ensembl (version 74) gene trees^28^ as described before^29,30^ and illustrated in Figure 1A and **1B**. Our definition of gene age tries to date the last time a gene was involved in a gene birth event. In the case of genes that have been duplicated (according to the Ensembl trees, Figure 1B), we consider the node of their most recent duplication - accordingly, daughter genes will have the same age (if none of them are involved in more recent duplication). In the case of singleton genes (i.e., those without detectable duplicates), we consider their gene age as the last common ancestor of all species that have orthologs (Figure 1A). In other words, the gene age is the gene family birth event (see review by Capra and coworkers^20^ about different ways to date gene ages).

The age groups Simiiformes, Catarrhini, Hominoidea, Hominidae, HomoPanGorilla and Homo sapiens were collapsed into Primate genes; while Rodent-specific merges Glires, Rodentia, Sciurognathi, Murinae and Mus musculus (here, we decided to name this group as Rodent because only the birth of 14 mouse genes were dated as Glires). Ten of the age groups correspond to the same ancestors in humans and mice (from FungiMetazoa to Eutheria) while the youngest group in each species correspond to gene births that occurred in Primates (in humans) or Rodents (in mice).

The gene ages for human and mouse genes are available at https://github.com/ricolab/TAD_Evolution and https://doi.org/10.5281/zenodo.10067329 (**Key resources table**).

#### TAD maps used in age co-localisation analysis

For the gene age co-localisation analysis (see **Quantification and statistical analysis**) we used the TAD genomic coordinates as defined by the original papers: hESCs,^31^ mESCs, ^38^ human GM12878 B-cell derived lymphoblastoid cells (original TADs^36^ and TADs from Gong and co-workers^43^), monocytes, neutrophils and naive CD4 T-cells (**Key resources Table**).^37^ We only modified the original data from Rao *et al* to remove the nested TADs, as they would have given duplicated co-localisations. All the TADs used are available at are available at https://github.com/ricolab/TAD_Evolution and https://doi.org/10.5281/zenodo.10067329 (**Key resources table**).

#### Gene essentiality data

Gene essentiality data from humans and mice was retrieved from OGEE database^44^ (**Key resources Table**).

#### Promoter-capture HiC (PCHi-C) networks

Networks were constructed with igraph^58^ R package from PCHiC data from mouse^45^ and human cells^37^ (**Key resources table**) by considering genomic regions as nodes and physical interactions as edges.^46^ We annotated each node with the genes and their ages. We created a subnetwork of young genes (Figure 1C), keeping interactions where at least one node contains a Primate or Rodent gene. Then, we grouped the nodes by gene and removed any duplicated edges so that each gene-gene interaction would only be represented once.

### QUANTIFICATION AND STATISTICAL ANALYSIS

#### Gene age co-localisation analysis

We generated the p-value matrices by performing 100 randomisations to obtain the variance of the distribution of the probabilities to find each pair of gene ages within the same TAD (see **Figure S1**). Then, using the normal distribution, the p-value was calculated as the probability of obtaining by chance alone the observed probability in the random distribution. We took as average the value of the null hypothesis, which was considered to be the probability of matching every age-pair without taking any TAD structure at all (considering the ages as being taken from a global pool rather than from any particular TAD). To facilitate the visualisation of these p-values, we calculated the natural cologarithm using a maximum value of 30. In order to differentiate between probabilities higher or lower than expected, the natural cologarithm (always positive since it is from a value lower than 1) was left unchanged (i.e., positive) if the probability of finding genes of an age pair together was higher than expected, or its sign was changed (becoming negative) for probabilities of finding a pair of ages lower than expected. The code to implement this is in function *getPairData*, available as part of the script *co-localisations_functions.R* at https://github.com/ricolab/TAD_Evolution and https://doi.org/10.5281/zenodo.10067329 (**Key resources table**).

The same approach was used to do the co-localisation analysis with fixed-size windows instead of TADs. The Euclidean distance between matrices was calculated with the dist() function in R.

#### TAD insulation

The insulation of TADs in GM12878 cells was calculated by averaging the two insulation scores of their boundaries, as calculated by Gong and co-workers.^43^ Then, TADs were grouped into five groups of similar size based on their TAD insulation scores, from low to high insulation. The original TAD boundary insulation scores were kindly provided by Aristotelis Tsirigos and we made them publicly available with his permission at https://github.com/ricolab/TAD_Evolution and https://doi.org/10.5281/zenodo.10067329 (**Key resources table**).

#### TAD stability across cell types

The stability of a TAD corresponds to the number of cell types where its boundaries are present. To study the stability of TADs across cell types, we used the TAD maps and associated stability scores calculated by McArthur and Capra^9^ (**Key resources table**). Their dataset includes definitions for 14,345 TAD boundaries and assigns to each of them a parameter, the stability percentile, that is associated with the number of cell types where it is present (which varies between one and 37). For example, if a TAD boundary is present in only one cell type, it corresponds to a stability percentile of 0.113175, but if it is present in 24 cell types, then its stability percentile is 0.901778. All TAD boundaries in this dataset span 100 kb.

We separated the dataset of TAD boundaries in two groups of approximately the same size depending on their presence in different cell types:

the “stable” group, which was made of TAD boundaries present in five or more different cell types, including 7,509 TAD boundaries (52%).
and the “unstable” group, which was made using the remaining 6,836 TAD boundaries (48%), present in less than five different cell types.

For each of these two groups, we created a list of TADs as the region between two consecutive TAD boundaries by taking the central value between the beginning and end of each TAD boundary definition. In addition, we removed all the TADs that were nesting a TAD boundary from the other group (so that the final TAD definitions in either group were empty of other TAD boundaries). Using these criteria, we obtained two datasets:

one with 4749 stable TADs, with an average length of 154 kb.
another one with 4086 unstable TADs, with an average length of 268 kb.

For each of these two TAD groups, we calculated the percentage of old genes using our dataset of gene ages (taking into account our definition of “old gene”, Figure 1C), defining as “old-depleted TADs” those that contained less than 10% of old genes. We observed that for the stable TADs, 323 out of 3031 had less than 10% of old genes (so the ratio corresponds to 0.1066), while for unstable TADs this filter is passed by 284 out of 1795 (ratio: 0.1582). This displays a higher amount of young TADs in unstable TADs. To assess the significance of this difference, we repeated the analysis 100 times for each group after randomising the gene age labels. We observed that all the randomised cases fall in between these values (see **Figure S6**), therefore showing that this difference is indeed statistically significant. The data and code to reproduce the analysis is available at https://github.com/ricolab/TAD_Evolution and https://doi.org/10.5281/zenodo.10067329.

#### Enrichment of essential genes

The significance of the enrichments of essential genes in the different sub-groups of genes was calculated with Chi-squared tests using the chi.square() function in R, and the significance threshold was set at p < 0.05. The code to reproduce the analysis is available at https://github.com/ricolab/TAD_Evolution and https://doi.org/10.5281/zenodo.10067329.

## References

1. Rada-Iglesias, A., Grosveld, F.G., and Papantonis, A. (2018). Forces driving the three-dimensional folding of eukaryotic genomes. Mol. Syst. Biol. 14, e8214.

2. Stadhouders, R., Filion, G.J., and Graf, T. (2019). Transcription factors and 3D genome conformation in cell-fate decisions. Nature 569, 345–354.

3. Krefting, J., Andrade-Navarro, M.A., and Ibn-Salem, J. (2018). Evolutionary stability of topologically associating domains is associated with conserved gene regulation. BMC Biol. 16, 87.

4. Lazar, N.H., Nevonen, K.A., O’Connell, B., McCann, C., O’Neill, R.J., Green, R.E., Meyer, T.J., Okhovat, M., and Carbone, L. (2018). Epigenetic maintenance of topological domains in the highly rearranged gibbon genome. Genome Res. 28, 983– 997.

5. Liao, Y., Zhang, X., Chakraborty, M., and Emerson, J.J. (2021). Topologically associating domains and their role in the evolution of genome structure and function in. Genome Res. 31, 397–410.

6. Ibn-Salem, J., Muro, E.M., and Andrade-Navarro, M.A. (2017). Co-regulation of paralog genes in the three-dimensional chromatin architecture. Nucleic Acids Res. 45, 81–91.

7. Long, H.S., Greenaway, S., Powell, G., Mallon, A.-M., Lindgren, C.M., and Simon, M.M. (2022). Making sense of the linear genome, gene function and TADs. Epigenetics Chromatin 15, 4.

8. Harmston, N., Ing-Simmons, E., Tan, G., Perry, M., Merkenschlager, M., and Lenhard, B. (2017). Topologically associating domains are ancient features that coincide with Metazoan clusters of extreme noncoding conservation. Nat. Commun. 8, 441.

9. McArthur, E., and Capra, J.A. (2021). Topologically associating domain boundaries that are stable across diverse cell types are evolutionarily constrained and enriched for heritability. Am. J. Hum. Genet. 108, 269–283.

10. Fudenberg, G., and Pollard, K.S. (2019). Chromatin features constrain structural variation across evolutionary timescales. Proc. Natl. Acad. Sci. U. S. A. 116, 2175– 2180.

11. Huynh, L., and Hormozdiari, F. (2019). TAD fusion score: discovery and ranking the contribution of deletions to genome structure. Genome Biol. 20, 60.

12. Whalen, S., and Pollard, K.S. (2019). Most chromatin interactions are not in linkage disequilibrium. Genome Res. 29, 334–343.

13. Canela, A., Maman, Y., Jung, S., Wong, N., Callen, E., Day, A., Kieffer-Kwon, K.-R., Pekowska, A., Zhang, H., Rao, S.S.P., et al. (2017). Genome Organization Drives Chromosome Fragility. Cell 170, 507–521.e18.

14. Farré, M., Robinson, T.J., and Ruiz-Herrera, A. (2015). An Integrative Breakage Model of genome architecture, reshuffling and evolution: The Integrative Breakage Model of genome evolution, a novel multidisciplinary hypothesis for the study of genome plasticity. Bioessays 37, 479–488.

15. Capra, J.A., Stolzer, M., Durand, D., and Pollard, K.S. (2013). How old is my gene? Trends Genet. 29, 659–668.

16. Domazet-Lošo, T., and Tautz, D. (2010). A phylogenetically based transcriptome age index mirrors ontogenetic divergence patterns. Nature 468, 815–818.

17. Chen, W.-H., Trachana, K., Lercher, M.J., and Bork, P. (2012). Younger genes are less likely to be essential than older genes, and duplicates are less likely to be essential than singletons of the same age. Mol. Biol. Evol. 29, 1703–1706.

18. Rancati, G., Moffat, J., Typas, A., and Pavelka, N. (2018). Emerging and evolving concepts in gene essentiality. Nat. Rev. Genet. 19, 34–49.

19. Bartha, I., di Iulio, J., Venter, J.C., and Telenti, A. (2018). Human gene essentiality. Nat. Rev. Genet. 19, 51–62.

20. Capra, J.A., Stolzer, M., Durand, D., and Pollard, K.S. (2013). How old is my gene? Trends Genet. 29, 659–668.

21. Juan, D., Rico, D., Marques-Bonet, T., Fernández-Capetillo, O., and Valencia, A. (2014). Late-replicating CNVs as a source of new genes. Biol. Open 3, 231.

22. Van de Peer, Y., Mizrachi, E., and Marchal, K. (2017). The evolutionary significance of polyploidy. Nat. Rev. Genet. 18, 411–424.

23. Van de Peer, Y., Maere, S., and Meyer, A. (2009). The evolutionary significance of ancient genome duplications. Nat. Rev. Genet. 10, 725–732.

24. Makino, T., and McLysaght, A. (2010). Ohnologs in the human genome are dosage balanced and frequently associated with disease. Proc. Natl. Acad. Sci. U. S. A. 107, 9270–9274.

25. Sacerdot, C., Louis, A., Bon, C., Berthelot, C., and Roest Crollius, H. (2018). Chromosome evolution at the origin of the ancestral vertebrate genome. Genome Biol. 19, 166.

26. Ohno, S. (1970). Evolution by Gene Duplication. 10.1007/978-3-642-86659-3.

27. Marlétaz, F., Firbas, P.N., Maeso, I., Tena, J.J., Bogdanovic, O., Perry, M., Wyatt, C.D.R., de la Calle-Mustienes, E., Bertrand, S., Burguera, D., et al. (2018). Amphioxus functional genomics and the origins of vertebrate gene regulation. Nature 564, 64–70.

28. Herrero, J., Muffato, M., Beal, K., Fitzgerald, S., Gordon, L., Pignatelli, M., Vilella, A.J., Searle, S.M.J., Amode, R., Brent, S., et al. (2016). Ensembl comparative genomics resources. Database 2016. 10.1093/database/bav096.

29. Juan, D., Rico, D., Marques-Bonet, T., Fernández-Capetillo, O., and Valencia, A. (2013). Late-replicating CNVs as a source of new genes. Biol. Open 2, 1402–1411.

30. Rigau, M., Juan, D., Valencia, A., and Rico, D. (2019). Intronic CNVs and gene expression variation in human populations. PLoS Genet. 15, e1007902.

31. Ma, W., Ay, F., Lee, C., Gulsoy, G., Deng, X., Cook, S., Hesson, J., Cavanaugh, C., Ware, C.B., Krumm, A., et al. (2015). Fine-scale chromatin interaction maps reveal the cis-regulatory landscape of human lincRNA genes. Nat. Methods 12, 71–78.

32. Martello, G., and Smith, A. (2014). The nature of embryonic stem cells. Annu. Rev. Cell Dev. Biol. 30, 647–675.

33. Chakraborty, C., and Agoramoorthy, G. (2012). Stem cells in the light of evolution. Indian J. Med. Res. 135, 813–819.

34. Agata, K., Nakajima, E., Funayama, N., Shibata, N., Saito, Y., and Umesono, Y. (2006). Two different evolutionary origins of stem cell systems and their molecular basis. Semin. Cell Dev. Biol. 17, 503–509.

35. Zufferey, M., Tavernari, D., Oricchio, E., and Ciriello, G. (2018). Comparison of computational methods for the identification of topologically associating domains. Genome Biol. 19, 217.

36. Rao, S.S.P., Huntley, M.H., Durand, N.C., Stamenova, E.K., Bochkov, I.D., Robinson, J.T., Sanborn, A.L., Machol, I., Omer, A.D., Lander, E.S., et al. (2014). A 3D map of the human genome at kilobase resolution reveals principles of chromatin looping. Cell 159, 1665–1680.

37. Javierre, B.M., Burren, O.S., Wilder, S.P., Kreuzhuber, R., Hill, S.M., Sewitz, S., Cairns, J., Wingett, S.W., Várnai, C., Thiecke, M.J., et al. (2016). Lineage-Specific Genome Architecture Links Enhancers and Non-coding Disease Variants to Target Gene Promoters. Cell 167, 1369–1384.e19.

38. Bonev, B., Mendelson Cohen, N., Szabo, Q., Fritsch, L., Papadopoulos, G.L., Lubling, Y., Xu, X., Lv, X., Hugnot, J.-P., Tanay, A., et al. (2017). Multiscale 3D Genome Rewiring during Mouse Neural Development. Cell 171, 557–572.e24.

39. Chang, L.-H., Ghosh, S., and Noordermeer, D. (2020). TADs and Their Borders: Free Movement or Building a Wall? J. Mol. Biol. 432, 643–652.

40. Dixon, J.R., Selvaraj, S., Yue, F., Kim, A., Li, Y., Shen, Y., Hu, M., Liu, J.S., and Ren, B. (2012). Topological domains in mammalian genomes identified by analysis of chromatin interactions. Nature 485, 376–380.

41. Cubeñas-Potts, C., and Corces, V.G. (2015). Topologically Associating Domains: An invariant framework or a dynamic scaffold? Nucleus 6, 430–434.

42. Allou, L., and Mundlos, S. (2023). Disruption of regulatory domains and novel transcripts as disease-causing mechanisms. Bioessays 45, e2300010.

43. Gong, Y., Lazaris, C., Sakellaropoulos, T., Lozano, A., Kambadur, P., Ntziachristos, P., Aifantis, I., and Tsirigos, A. (2018). Stratification of TAD boundaries reveals preferential insulation of super-enhancers by strong boundaries. Nat. Commun. 9, 542.

44. Chen, W.-H., Lu, G., Chen, X., Zhao, X.-M., and Bork, P. (2017). OGEE v2: an update of the online gene essentiality database with special focus on differentially essential genes in human cancer cell lines. Nucleic Acids Res. 45, D940–D944.

45. Schoenfelder, S., Furlan-Magaril, M., Mifsud, B., Tavares-Cadete, F., Sugar, R., Javierre, B.-M., Nagano, T., Katsman, Y., Sakthidevi, M., Wingett, S.W., et al. (2015). The pluripotent regulatory circuitry connecting promoters to their long-range interacting elements. Genome Res. 25, 582–597.

46. Pancaldi, V., Carrillo-de-Santa-Pau, E., Javierre, B.M., Juan, D., Fraser, P., Spivakov, M., Valencia, A., and Rico, D. (2016). Integrating epigenomic data and 3D genomic structure with a new measure of chromatin assortativity. Genome Biol. 17, 152.

47. Kalsoom, U.-E.-, Klopocki, E., Wasif, N., Tariq, M., Khan, S., Hecht, J., Krawitz, P., Mundlos, S., and Ahmad, W. (2013). Whole exome sequencing identified a novel zinc-finger gene ZNF141 associated with autosomal recessive postaxial polydactyly type A. J. Med. Genet. 50, 47–53.

48. Xie, T., Yang, Q.-Y., Wang, X.-T., McLysaght, A., and Zhang, H.-Y. (2016). Spatial Colocalization of Human Ohnolog Pairs Acts to Maintain Dosage-Balance. Mol. Biol. Evol. 33, 2368–2375.

49. Gaboriaud, J., and Wu, P.-Y.J. (2019). Insights into the Link between the Organization of DNA Replication and the Mutational Landscape. Genes 10. 10.3390/genes10040252.

50. Li, D., He, M., Tang, Q., Tian, S., Zhang, J., Li, Y., Wang, D., Jin, L., Ning, C., Zhu, W., et al. (2022). Comparative 3D genome architecture in vertebrates. BMC Biol. 20, 99.

51. Álvarez-González, L., Burden, F., Doddamani, D., Malinverni, R., Leach, E., Marín-García, C., Marín-Gual, L., Gubern, A., Vara, C., Paytuví-Gallart, A., et al. (2022). 3D chromatin remodelling in the germ line modulates genome evolutionary plasticity. Nat. Commun. 13, 2608.

52. Okhovat, M., VanCampen, J., Lima, A.C., Nevonen, K.A., Layman, C.E., Ward, S., Herrera, J., Stendahl, A.M., Yang, R., Harshman, L., et al. (2023). TAD Evolutionary and functional characterization reveals diversity in mammalian TAD boundary properties and function. bioRxiv. 10.1101/2023.03.07.531534.

53. Majic, P., and Payne, J.L. (2020). Enhancers Facilitate the Birth of De Novo Genes and Gene Integration into Regulatory Networks. Mol. Biol. Evol. 37, 1165–1178.

54. Carelli, F.N., Liechti, A., Halbert, J., Warnefors, M., and Kaessmann, H. (2018). Repurposing of promoters and enhancers during mammalian evolution. Nat. Commun. 9, 4066.

55. Davidson, B.S.A., Arcila-Galvis, J.E., Trevisan-Herraz, M., Mikulasova, A., Brackley, C.A., Russell, L.J., and Rico, D. (2023). The discovery of an evolutionarily conserved enhancer within the MYEOV locus suggests an unexpected role for this non-coding region in cancer. bioRxiv, 2023.09.18.558245. 10.1101/2023.09.18.558245.

56. Santiago-Algarra, D., Souaid, C., Singh, H., Dao, L.T.M., Hussain, S., Medina-Rivera, A., Ramirez-Navarro, L., Castro-Mondragon, J.A., Sadouni, N., Charbonnier, G., et al. (2021). Epromoters function as a hub to recruit key transcription factors required for the inflammatory response. Nat. Commun. 12, 6660.

57. Lewin, H.A., Richards, S., Lieberman Aiden, E., Allende, M.L., Archibald, J.M., Bálint, M., Barker, K.B., Baumgartner, B., Belov, K., Bertorelle, G., et al. (2022). The Earth BioGenome Project 2020: Starting the clock. Proc. Natl. Acad. Sci. U. S. A. 119. 10.1073/pnas.2115635118.

58. Csárdi, G., Nepusz, T., Müller, K., Horvát, S., Traag, V., Zanini, F., and Noom, D. (2023). igraph for R: R interface of the igraph library for graph theory and network analysis (Zenodo) 10.5281/ZENODO.7682609.

