## Supplemental Figures 1-6 for "Evolutionary analysis of gene ages across TADs associates chromatin topology with whole genome duplications"

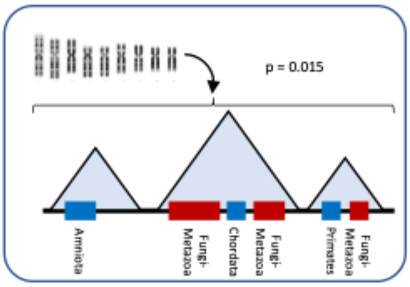


1. Calculate the probability of finding a Fungi-Metazoa gene in the same TAD where there is already another Fungi-Metazea gene

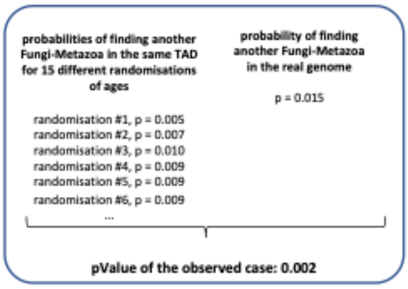


2. Compare the observed probability to the probabilities of 100 different randomisations where the ages have been scrambled within the table assigning ages to genes.

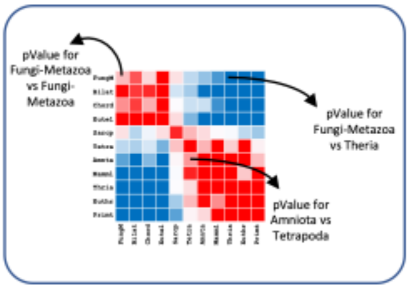


3. Repeat this process for every age combination (the matrix is not exactly symmetrical due to statistical variance). We observe here the two-blocks pattern, one for old genes, the other for young genes (see **Figure 2B** in the main text).

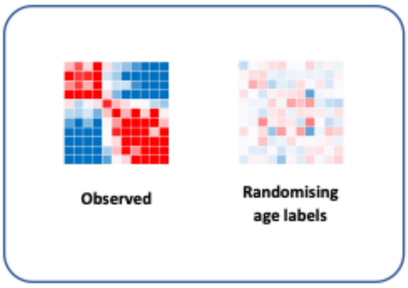


4. To understand the statistical fluctuations of the randomisations, we compare the observed case to the result after randomising the labels of also the initial case (so we are comparing a randomisation against the other 100 randomisations). As expected, the pValues of the second case are well above 5%, and any pattern is broken.

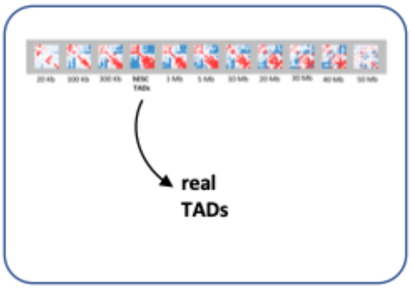


5. Repeat the process using several sizes of regular windows and compare the result with the real TAD definitions (see **Figure 3** in the main text).

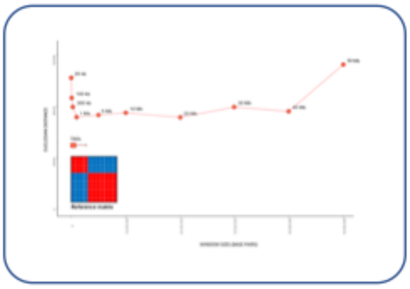

6. Compare each of those cases to a hypothetical two-block matrix using the Euclidean distance.

**Figure S1. Schematic overview of our statistical approaches to calculate gene age co-localisations** (related to **Figures 2** and **3**). See STAR Methods for details.

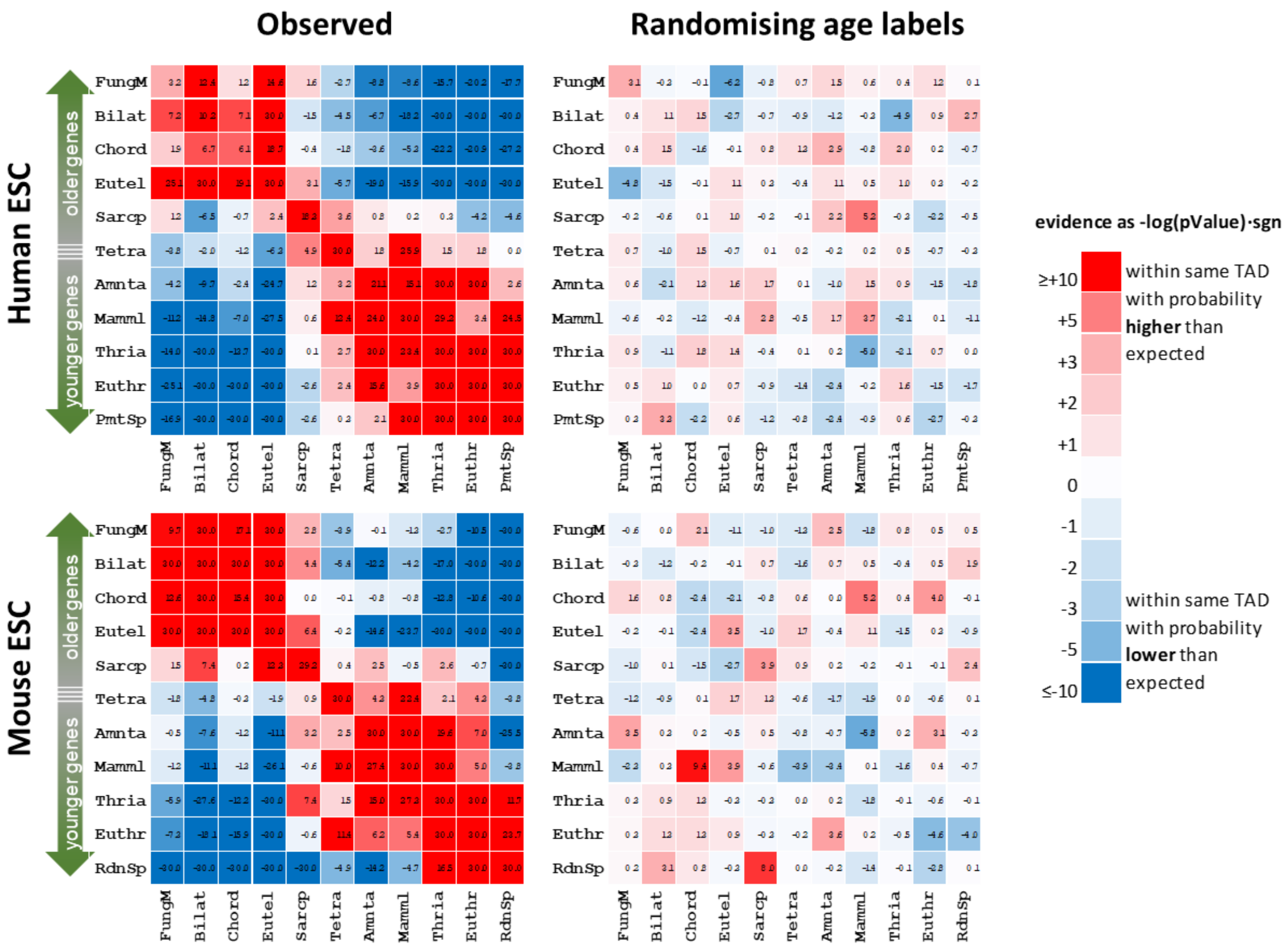


**Figure S2. Gene age colocalisation results with TADs from hESCs and mESCs** (related to **Figure 2**). **Left:** observed p-values (same as **Figure 2B,** including numerical values). **Right:** heatmaps that show that the observed colocalisation patterns are lost even with just five randomisations. See **Figure 1C** for the correspondence between abbreviated and full gene ages, and **Key resources table** for the sources of TADs.

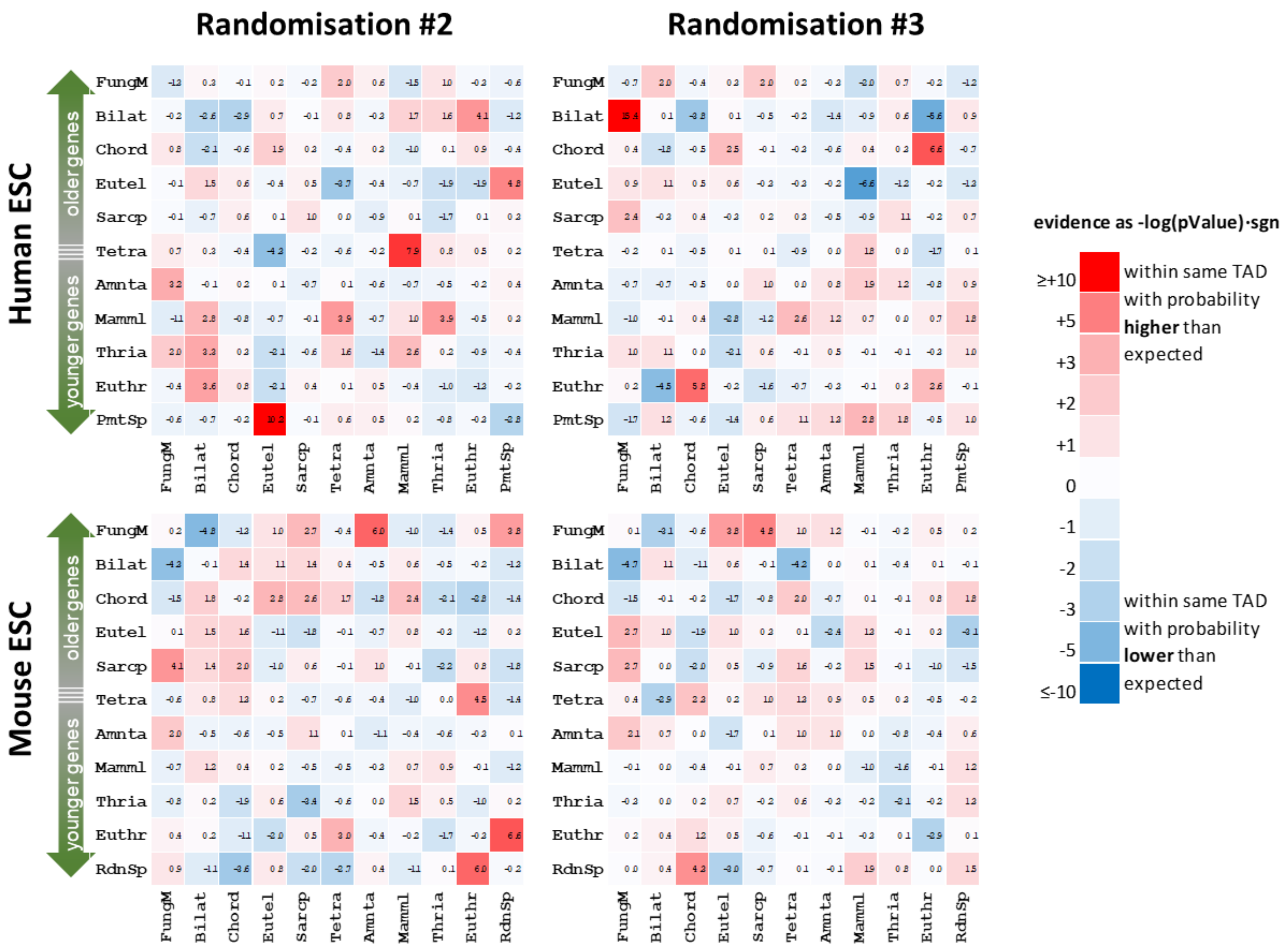


**Figure S3. Two additional randomisations with different seeds** (related to **Figure 2B**). Sample displaying two further randomisations (calculated the same way as in **Figure S2** (right heatmaps) for human and mouse ESCs, using two different seeds with five randomisations each. The lack of clear patterns in the colocalisation of coetaneous genes becomes apparent again. See **Figure 1C** for the correspondence between abbreviated and full gene ages

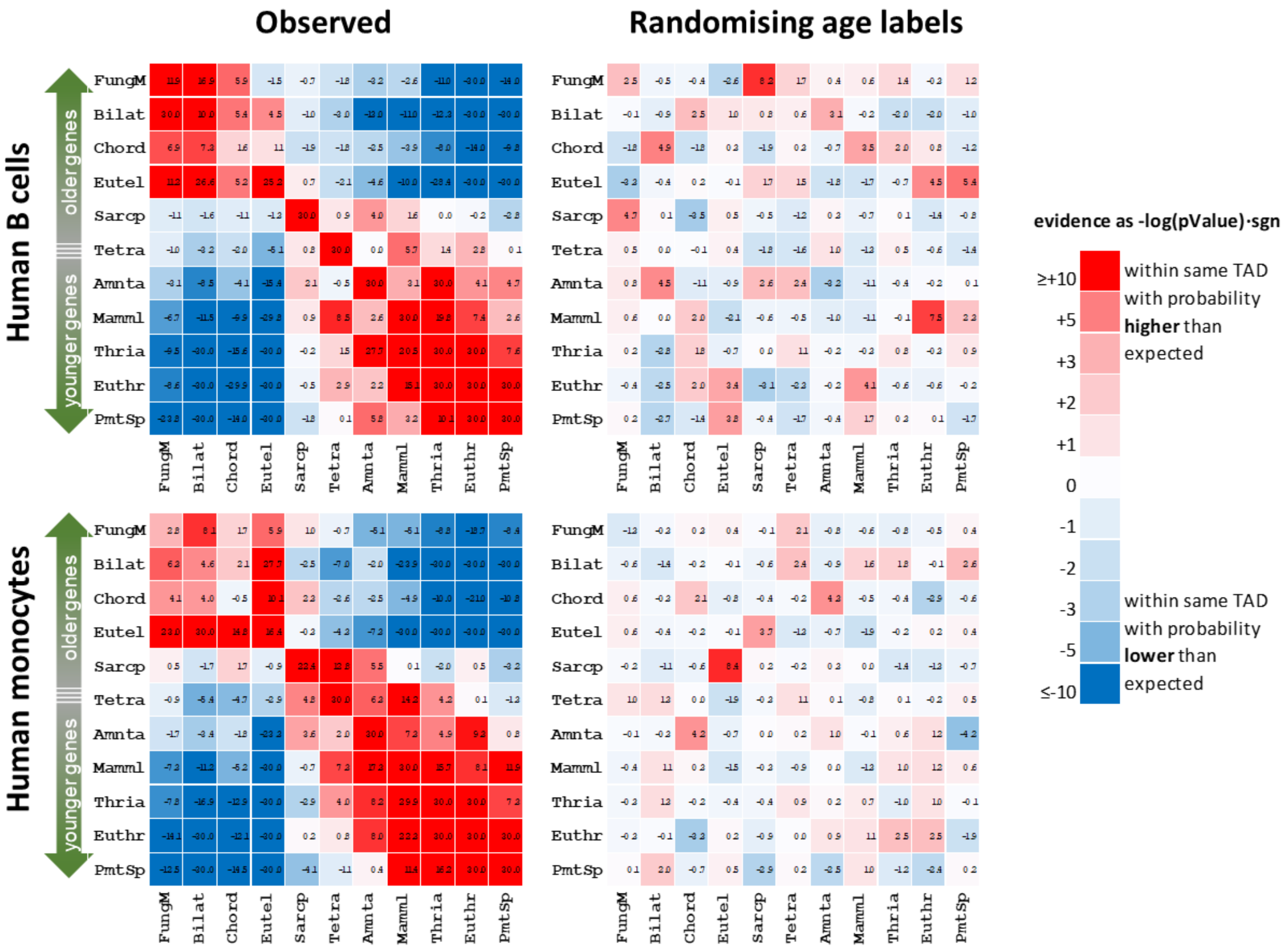


**Figure S4. Gene age colocalisation analysis with TADs from human monocytes and lymphoblastoid B-cells** (related to **Figure 2B**)**.** Same analysis as in **Figure S2**, related to **Figure 2B** but using TADs of human B-derived lymphoblastoid cells (GM12878) and monocytes (**Key resources table**) with five randomisations. For further details see **Figure 2**. See **Figure 1C** for the correspondence between abbreviated and full gene ages.

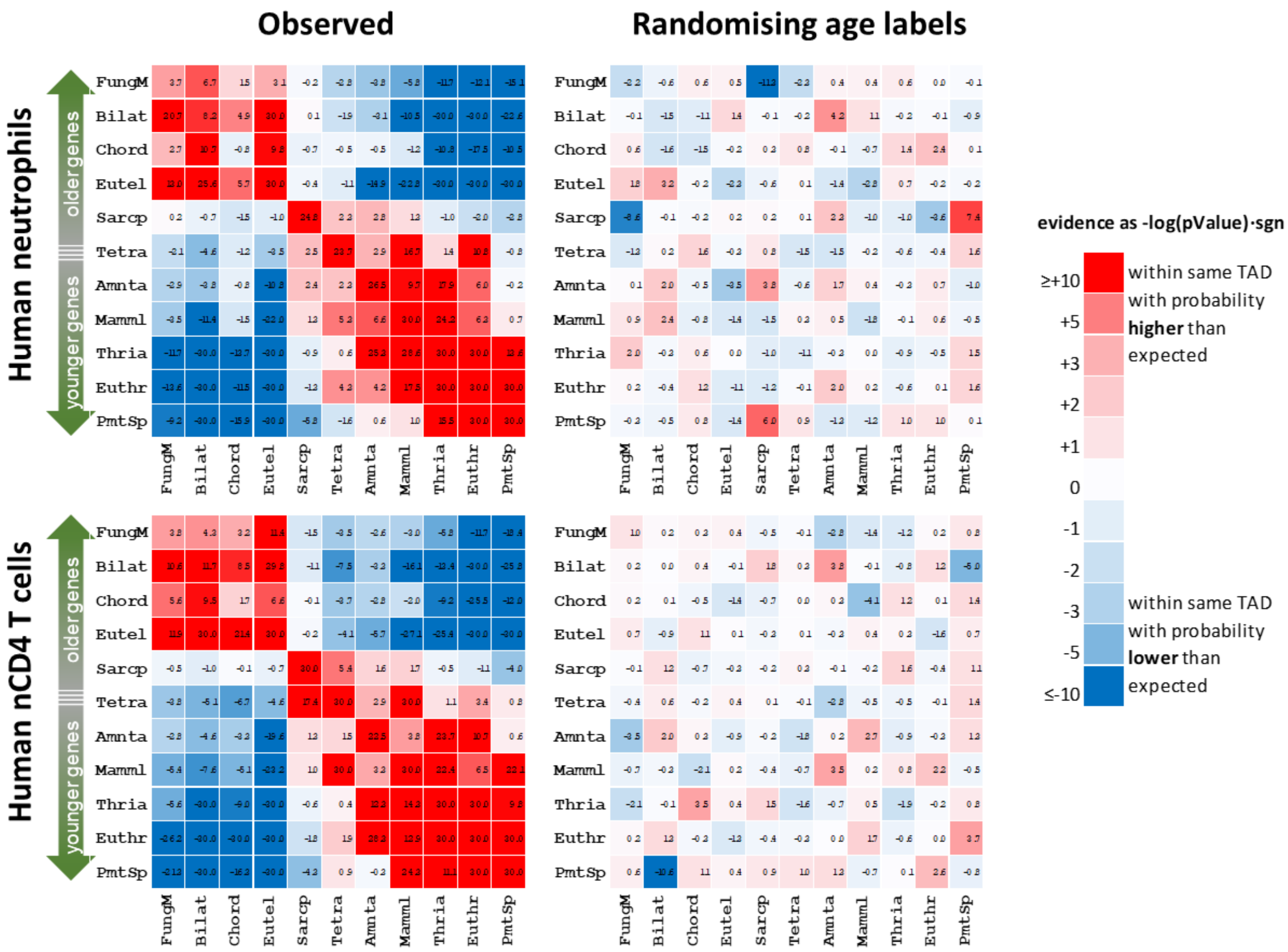


**Figure S5. Gene age colocalisation analysis with TADs from human neutrophils and T-cells** (related to **Figure 2B**). Same analysis as in **Figure S2**, but using TADs of human neutrophils and naive CD4 T cells with five randomisations(**Key resources table**). For further details see **Figure 2B**. See **Figure 1B** for the correspondence between abbreviated and full gene ages.

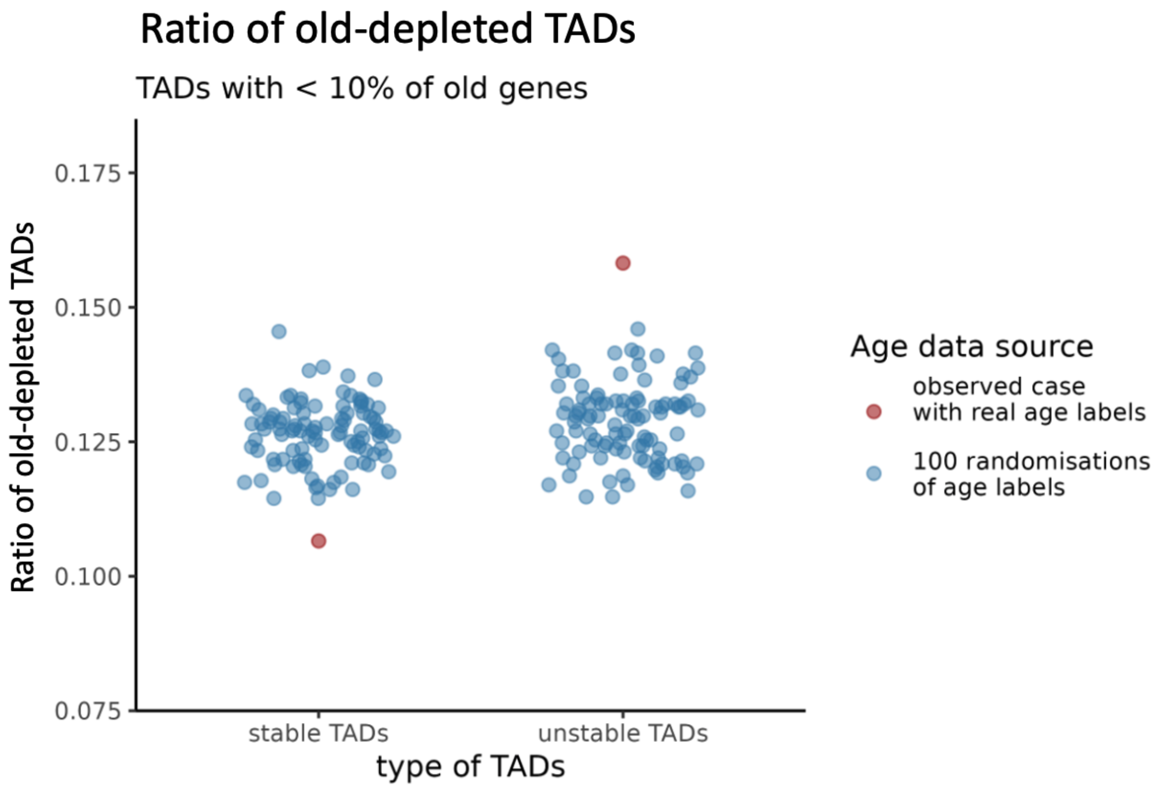


**Figure S6. TADs depleted in old genes are unstable across cell types** (related to **Figure 4**)**.** Using the integrated TAD maps across 37 human cell types from McArthur and Capra (**Key resources table**), we classified TADs into two groups of similar size: those stable across cell types (with boundaries present in 5 to 37 cell types) and those unstable (boundaries present in 4 or fewer cell types). In each group, we calculated the ratio of old-depleted TADs, defined as those TADs with less than 10% young (Primate) ages. The y-axis shows the observed ratio of old-depleted TADs in each group of TADs (red points). We repeated this calculation 100 times after randomly assigning the age labels (blue points).
